# In vivo repair of a protein underlying a neurological disorder by programmable RNA editing

**DOI:** 10.1101/2020.02.26.966820

**Authors:** John R Sinnamon, Susan Y Kim, Jenna R Fisk, Zhen Song, Hiroyuki Nakai, Sophia Jeng, Shannon K McWeeney, Gail Mandel

## Abstract

RNA base editing is gaining momentum as an approach to repair mutations, but its application to neurological disease has not been established. We have succeeded in directed transcript editing of a pathological mutation in a mouse model of the neurodevelopmental disease, Rett syndrome. Specifically, we directed editing of a guanosine to adenosine mutation in RNA encoding Methyl CpG Binding Protein 2 (MECP2). Repair was mediated by injecting the hippocampus of juvenile Rett mice with an adeno-associated virus expressing both an engineered enzyme containing the catalytic domain of Adenosine Deaminase Acting on RNA 2 and a *Mecp2* targeting guide. After one month, 50% of *Mecp2* RNA was recoded in three different hippocampal neuronal subtypes, and the ability of MeCP2 protein to associate with heterochromatin was similarly restored to 50% of wild-type levels. This study represents the first *in vivo* programmable RNA editing applied to a model of neurological disease.

## Introduction

Adenosine deaminase enzymes catalyze a post-transcriptional adenosine to inosine exchange in RNA (A-to-I RNA editing) [1-5]. Inosine is treated as guanosine by the translational machinery [6], resulting in codon changes with consequences for protein function [7-9]. Two catalytically active Adenosine Deaminase Acting on RNA enzymes, ADAR1 and 2, are expressed to high levels in the CNS [10]. The native proteins are recruited to their natural target RNAs, which can be overlapping or unique to each molecule [3,9,11,12], by intrinsic domains that recognize double stranded RNA [3,5]. Several pioneering studies have paved the way for programmable A-to-I editing of exogenous RNAs by harnessing the deaminase domains of ADAR1 or 2 [13-15]. Many of these studies utilized fusions of the ADAR deaminase domain with a heterologous RNA binding protein, for example Cas13 or the bacteriophage λN peptide, with a RNA guide specific for the target RNA substrate [13,15-21]. Endogenous RNAs have been shown to be repaired by programmable RNA editing in cell lines and primary cells [17-19,22]. A recent study demonstrated repair in mouse models of Duchenne muscular dystrophy and ornithine transcarbamylase deficiency [21], establishing the utility of this approach for both muscle and liver. However, both of these tissues have the advantage of relative cellular homogeneity. The large cellular complexity in the CNS raises the question of whether RNA structure and RNA binding proteins in different cell types will create a more or less accessible transcript for RNA repair by programmable editing. Therefore, direct tests in a mouse model of neurological disease are clearly warranted.

Mouse models of Rett syndrome are ideally suited to test the efficacy of programmable RNA editing *in vivo*. Rett syndrome is caused by *de novo* loss-of-function mutations in the gene encoding the *X*-linked transcriptional regulator, MECP2 [23]. MeCP2 pathological mutations result in primarily a neurological phenotype characterized, in females, by regression of speech and purposeful hand motions, and the appearance of seizures, and respiratory abnormalities [24]. MeCP2 is expressed in most, if not all, neurons and glia [25-29], and previous studies have indicated that these cells can be targeted throughout the brain and spinal cord by adeno-associated virus 9 (AAV9) for facile analyses [30-34]. Importantly, a database of mutations causing Rett syndrome [35] indicates that 36% are caused by G>A mutations or C>T mutations that create opal stop codons, raising the possibility that adenosine deamination in these contexts may repair MeCP2 protein function. We previously generated a mouse line in which the mouse *Mecp2* gene contained the human patient mutation, *MECP2*^*317G>A*^ (R106Q) [19]. This mutation, resulting in classic Rett syndrome, is located in the DNA binding domain. Consequently, MeCP2 protein becomes unstable and has greatly reduced ability to bind to chromatin [19,36,37], easily quantifiable metrics for recovery of protein function. Indeed, by expressing the ADAR2 deaminase domain fused to the bacteriophage λN peptide (Editase) [15], together with a mouse *Mecp*2 guide containing the RNA sequence recognized by the λN peptide, endogenous *Mecp2*^*317G>A*^ RNA in cultured neurons was repaired efficiently [19]. However, whether we could virally deliver sufficient amounts of Editase *in vivo*, whether different neuronal types would be equally accessible to directed RNA editing, and the extent of off-target editing, remained open questions.

Here, using sequencing and imaging analyses, we test whether RNA can be repaired by programmable RNA editing in the nervous system. Using MeCP2 as a model we determine whether *Mecp2* RNA, in distinct neuronal subpopulations in postnatal brain, is accessible to programmable RNA repair, and whether MeCP2 protein function is also restored. We also determine the on- and off-target editing landscape in one of these populations of neurons by whole transcriptomic analysis.

## Results

### On-and off-target editing analyses of Mecp2 RNA

Six copies of a *Mecp2* targeting guide or a non-targeting guide (Supplementary Table S1) were cloned downstream of the U6 small nuclear RNA polymerase III promoter and upstream of the recombinant phage λN-hADAR2 deaminase domain protein (Editase). Because the target adenosine in *Mecp2*^*317G>A*^ is not in an ideal sequence context for ADAR2 editing [38], we utilized a version of Editase that contains a mutation, E488Q, resulting in hyperactivity [19,39]. The non-targeting guide lacks both the BoxB RNA hairpin recognized by the λN RNA binding domain in Editase as well as any *Mecp2* sequences. The HA epitope-tagged Editase^E488Q^ was placed under control of the human *Synapsin I* promoter to direct expression to neurons after stereotactic injection of the viruses (Figure 1a). We produced AAV vectors expressing each component by packaging these expression cassettes into the AAV PHP.B capsid.

**Figure 1:**
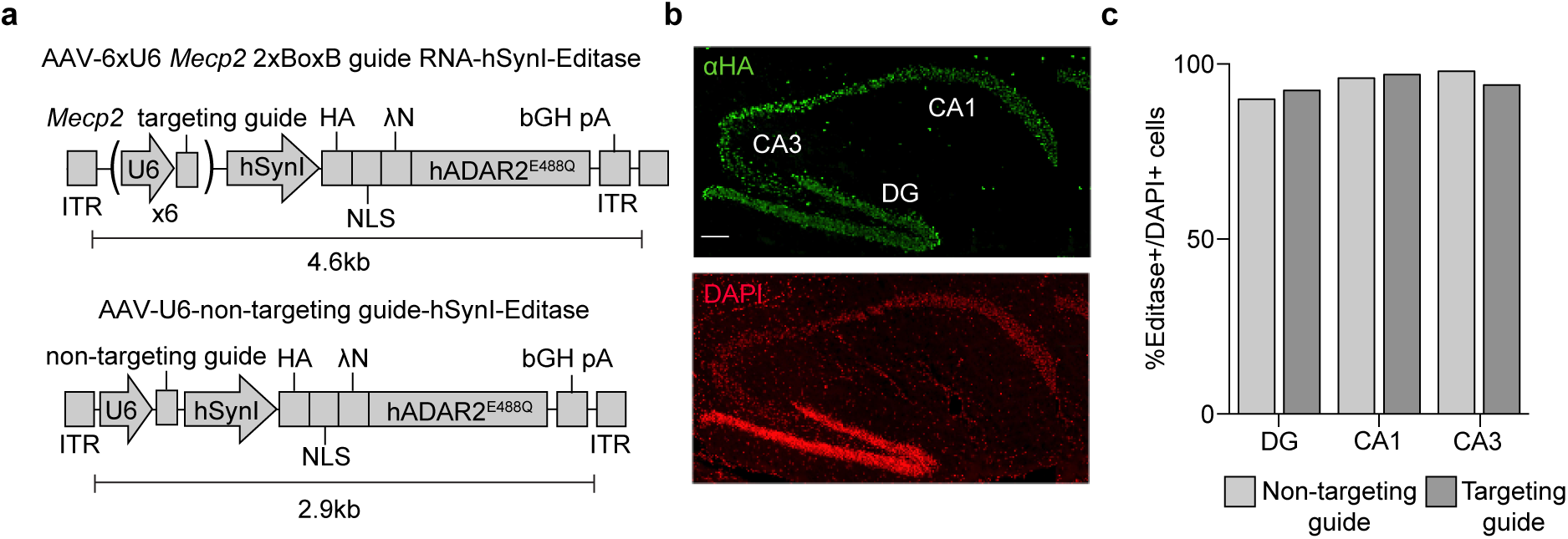
Hippocampal expression of RNA editing components following stereotaxic injection in the *Mecp2*^*317G>A*^ mouse model of Rett syndrome. a) Schematic showing the design of AAV Editase expression vectors. Each construct contains the human *Synapsin I* promoter for neuronal Editase expression and either (*top*) six copies of the U6 promoter each expressing a single copy of the *Mecp2* 2xBoxB targeting guide or a single copy of the human U6 promoter for expression of a small non-targeting RNA (*bottom*). b) Representative confocal images of a *Mecp2*^*317G>A*^ mouse three weeks after injection of AAV PHP.B vector into the hippocampus. Editase expression was detected by HA immuno-staining in the dentate gyrus (DG) and CA1 and CA3 pyramidal neuronal layers. Scale bar = 100 µm. c) Quantification of the percentage of HA-Editase positive cells for each guide virus relative to the total number of cells in each region (mean, n = 2 mice per condition). >100 cells were counted for each hippocampal region per replicate.

We first established expression of Editase by immunohistochemistry for the HA epitope tag after stereotaxic injection of both vectors into the hippocampus. We observed Editase expression from both vectors in the major neuronal populations of the hippocampus (Figure 1b, c). We then determined on- and off-target A-to-I editing rates within *Mecp2* by Sanger sequence analysis. RNA was isolated from the entire hippocampus as well as from different subpopulations of hippocampal neurons (CA1, CA3, dentate gyrus) isolated by laser-capture micro-dissection from each of three mice. In entire hippocampus we measured 35±7% A-to-I editing at the target adenosine with the *Mecp2* guide, with negligible editing observed with the non-targeting guide (Figure 2a). When cDNA was prepared from the individual hippocampal neuronal populations, the mean editing rates at the target site varied between 49-52% (DG, 49±9; CA1, 52±12; CA3, 49±7) and editing was again negligible with the non-targeting guide (Figure 2b). External to the guide region, no off-target sites were detected in *Mecp2* RNA. Within the guide region, we detected five bystander off-target editing sites, across all of the injected mice (Table 1). There was only one site with a consistently high rate of editing (18-65%). This site was detected in all of the mice regardless of neuronal population, and resulted in a silent mutation (E102E).

**Table 1:**
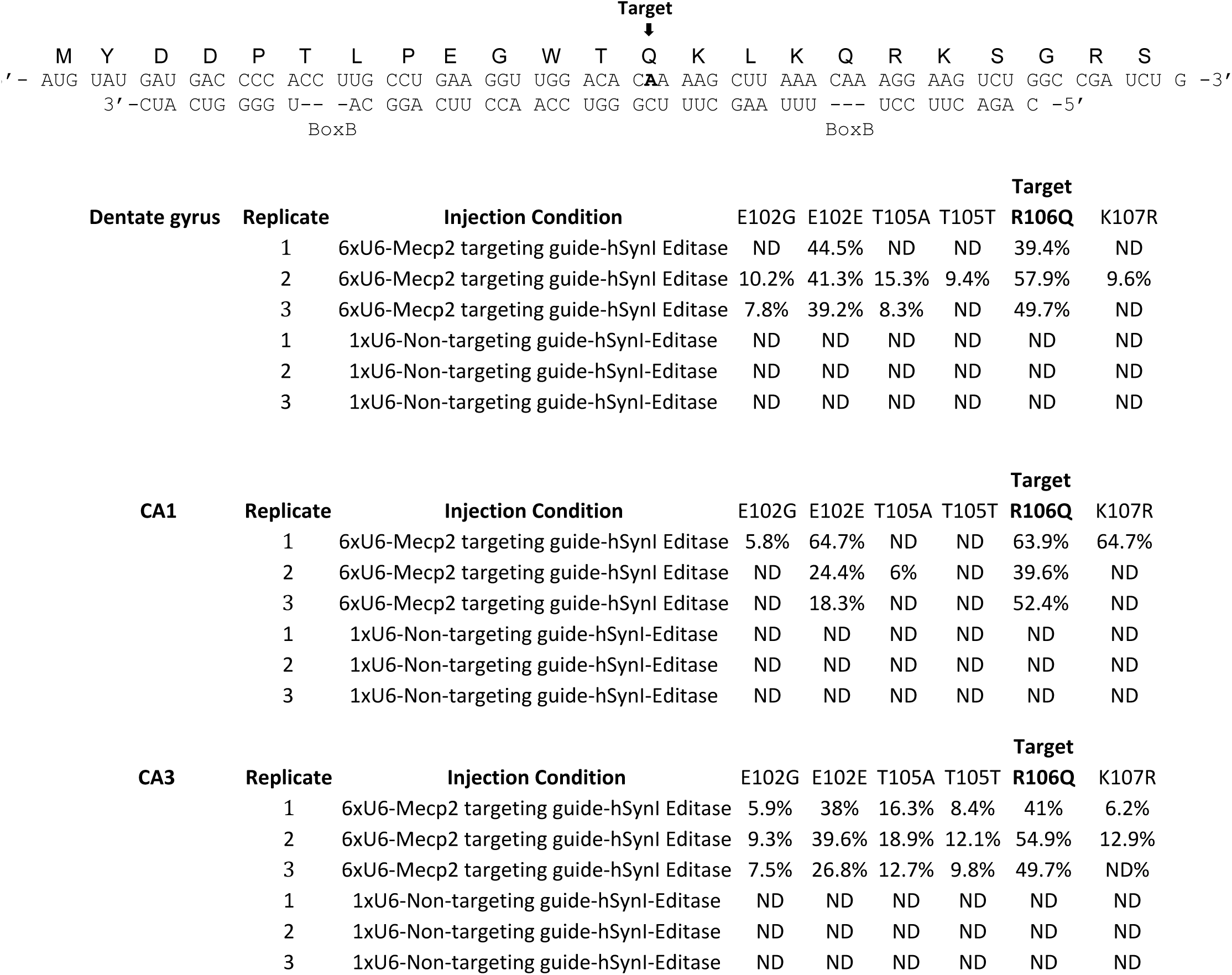
Editing of adenosines within *Mecp2* RNA identified by Sanger sequencing analysis. *Top, Mecp2* mRNA and deduced primary amino acid sequence (top row) relative to the guide RNA (bottom row). The target adenosine, *Mecp*^317^ in the RNA strand (MeCP2 R106Q) is bolded. *Bottom*, Rates of editing at the adenosines located within the guide region for each biological replicate. The detection limit for this assay was previously determined to be 5% editing [19]. Any sites that had ≤ 5% editing are listed as having no detectable editing (ND). There was no detectable editing in *Mecp2* RNA outside of the guide region.

**Figure 2:**
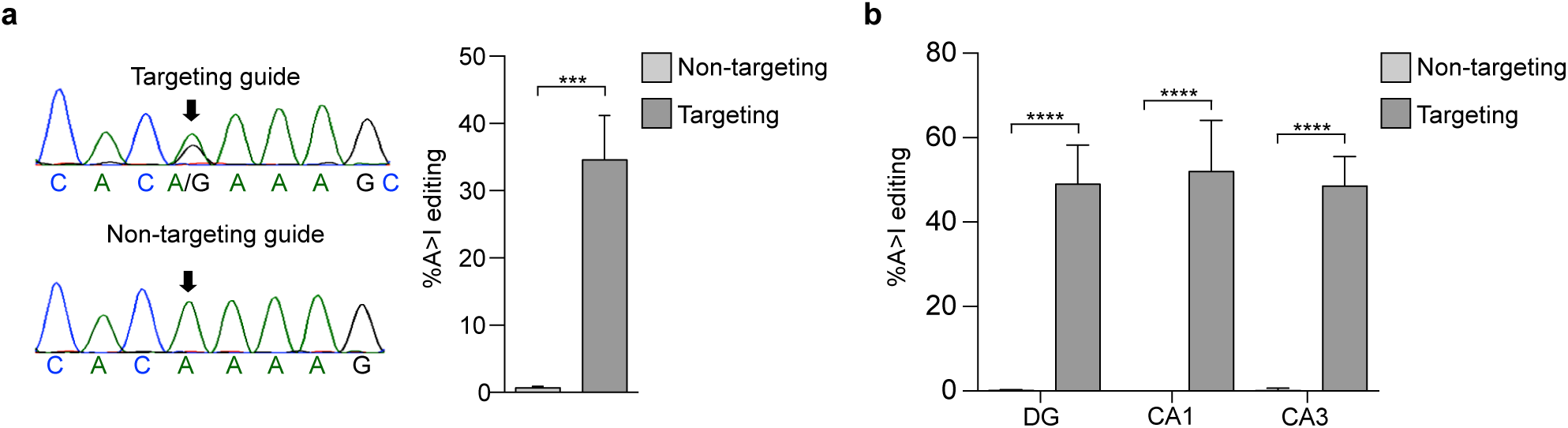
Efficient editing of *Mecp2* RNA following AAV injection directly into the hippocampus of P28 *Mecp2*^*317G>A*^ male mice. a) *Left*, sequencing chromatograms of cDNA from whole hippocampus, isolated three weeks after delivery of virus with Editase and indicated guides. Arrow denotes the on-target base. *Right*, quantification of editing (mean ± SD, n = 3 mice per condition)*** p<0.01 unpaired two-tailed *t*-test. b) Quantification of editing (mean ± SD, n = 3 mice per condition) by Sanger sequencing analysis of cDNA from indicated regions of the hippocampus isolated by laser capture micro-dissection three weeks after injection of virus. **** p < 0.001 one-way ANOVA and Tukey’s multiple comparison test.

We next performed a high throughput RNA-seq analysis on one of the neuronal populations, dentate gyrus, to determine A-to-I on-and off-target editing rates within the whole transcriptome (Figure 3). Whole-exomic sequencing was also performed on the same samples. Single nucleotide polymorphisms and edited sites in the non-injected control samples, presumably reflecting endogenous ADAR activity, were removed from the results presented herein. Importantly, the on-target and off-target editing rates within the guide region matched those from our Sanger sequencing analysis using the same RNA. Additionally, editing sites were not detected anywhere in the *Mecp2* RNA in the mice infected with the non-targeting guide virus. These results support our RNA seq bioinformatics pipeline.

**Figure 3:**
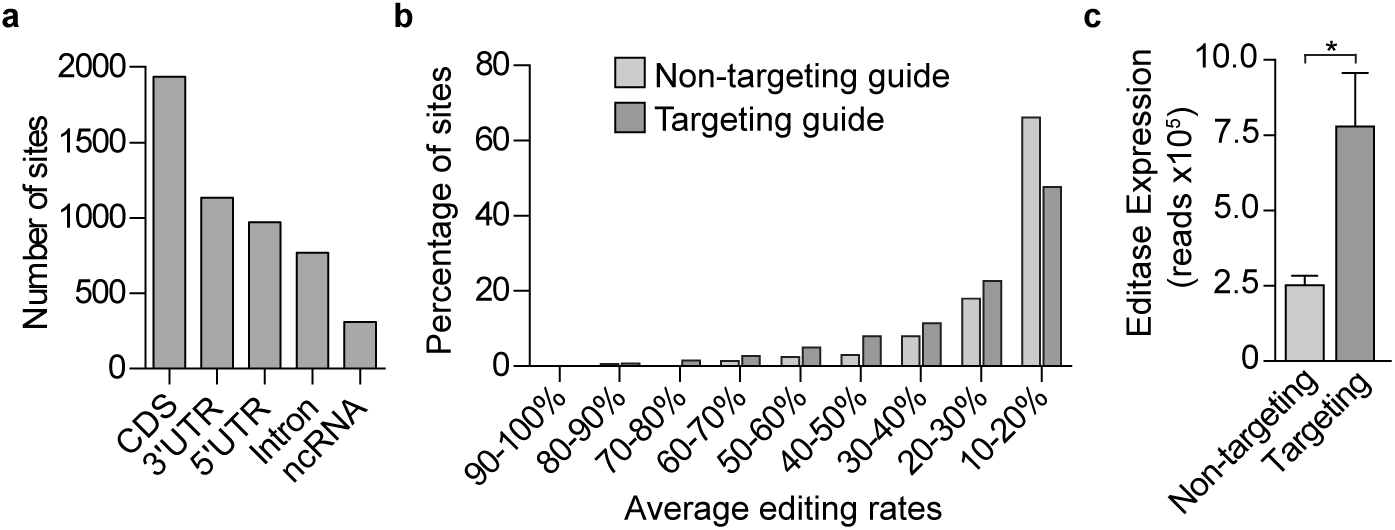
Off-target editing rates determined by whole transcriptomic analysis in the dentate gyrus is graded and dependent upon the level of Editase expression. a) Histogram showing the number of total off-target sites, independent of injection condition, located in coding sequences (CDS), 3’ untranslated regions (3’UTR), 5’ untranslated regions, introns and non-coding RNAs (ncRNA). b) Histogram showing the percentage of transcriptome wide RNA editing sites. Editing sites were binned according to the average editing rates across three biological replicates. c) Editase expression represented by the number of RNA-seq reads (mean ± SD, n = 3) that aligned to the Editase coding sequence for each injection condition. * p < 0.05, two tailed unpaired *t*-test.

### Global off target analysis

We identified 2984 off-target sites from the mice injected with the *Mecp2* targeting guide virus and 909 off-target sites from the mice injected with the non-targeting guide virus. The off-target sites in both conditions were distributed throughout the primary transcript (Figure 3a). When the number of sites and the percentages of A-to-I editing were considered, for both targeting and non-targeting guides, a majority (70% and 84%, respectively) represented editing at rates of ≤ 30% (Figure 3b). Consistent with a previous study [17], we found that nearly all (97%) of the off-target sites in the non-targeting condition were included in the sites in the *Mecp2* targeting condition, and that off-target editing was influenced by Editase levels (Figure 3b and 3c). To confirm that on-target editing of *Mecp2* RNA was guide-dependent and independent of Editase levels, we performed a new RNA-seq analysis on the same RNA from our previous neuronal culture study [19], where, fortuitously, the Editase level was higher in cells infected with the non-targeting virus (Supplementary Figure 1a). Despite the higher Editase levels in the non-targeting condition, there was no on-target editing without the *Mecp2* targeting guide (Supplementary Figure 1b). Further, off-target editing was influenced by Editase expression level (Supplementary Figure 1c). The U6 promoter number did not influence Editase protein levels in transfection analysis, suggesting that variability in Editase levels in neuronal cultures and *in vivo* reflect variability in viral infection parameters (Supplementary Figure 2).

### Repair of MeCP2 protein function

The *Mecp2*^*317G>A*^ mutation destabilizes MeCP2 protein *in vitro* and *in vivo*, resulting in greatly diminished binding to heterochromatin. We asked, at the single cell level, whether the broadly distributed off-target editing sites we detected would prevent MeCP2 protein functions, for example, by causing further destabilization, altering chromatin, or, for example, by preventing nuclear entry. Because MeCP2 binds to methylated DNA that is enriched in mouse satellite sequences in the pericentromeric heterochromatin, enrichment in heterochromatin has been an *in vivo* proxy for MeCP2 DNA binding ability [40,41]. Therefore, we used confocal microscopy to quantitate the enrichment of MeCP2 within regions of interest in neuronal heterochromatin foci in the hippocampus of two *Mecp2*^*317G>A*^ mice for each viral condition (see Methods). Two non-injected wild-type mice were used as controls. We analyzed the same fields-CA1, CA3 and dentate gyrus-that were used for Sanger sequencing analyses (Figure 2b). Neither the number of DAPI-labeled heterochromatin puncta per nucleus, nor the average size of heterochromatic puncta, was altered between virus infected and wild-type nuclei. In all fields, for CA3 and dentate gyrus sections, both mice infected with virus expressing the *Mecp2* targeting guide showed heterochromatic enrichment of MeCP2 protein (Figure 4a,d and c,f). Further, the amount of MeCP2 within heterochromatin relative to wild-type MeCP2, was nearly the same as the amount of on-target editing by Sanger and whole transcriptomic sequencing (compare Figure 4d,e and Figure 2b). In contrast, in all neurons infected with the non-targeting virus, MeCP2 protein was destabilized and not enriched in heterochromatin (Figure 4a-c).

**Figure 4:**
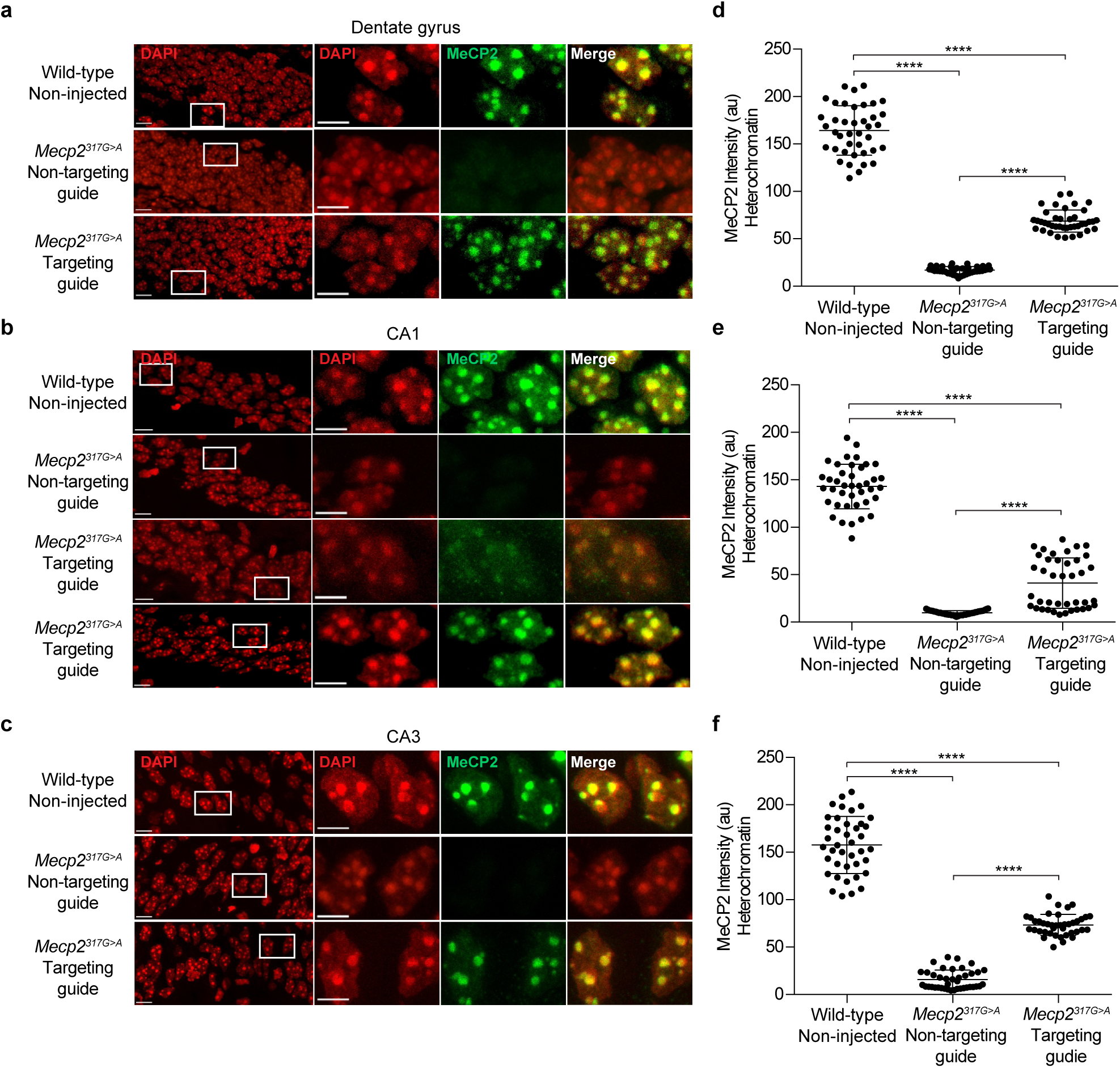
Programmable RNA editing restores the ability of MeCP2 to associate with heterochromatin in *Mecp2*^*317G>A*^ mice expressing Editase and *Mecp2* targeting guide.a-c) Confocal images of hippocampal neuronal nuclei immuno-labeled for MeCP2. DAPI staining outlines the nuclei and is enriched in the heterochromatic foci shown in the higher magnification images enclosed in the boxes. Scale bars, 10 µm, higher power; 5 µm lower power images. All images were acquired at the same intensity measurements. d-f) Quantification of immuno-labeled MeCP2 associated with heterochromatic foci (see methods). Each dot represents a single cell, 40 cells total from two mice. a and d, Dentate granule neurons; b and e, CA1 pyramidal neurons. c and f, CA3 pyramidal neurons. au, arbitrary units. **** p < 0.0001 by Kruskal-Wallis test and Dunn’s multiple comparisons test.

Interestingly, one mouse injected with the *Mecp2* targeting virus did not show MeCP2 enrichment within the heterochromatin of CA1 neurons, although Editase was expressed and enrichment was robust in both CA3 and dentate gyrus in the same mouse. This result is reflected in the bimodal distribution of heterochromatin-associated MeCP2 in the *Mecp2* targeting guide condition of CA1 neurons (Figure 4b,e). We cannot measure both RNA repair and immunohistochemistry in the same mouse for technical reasons. However, the lack of functional MeCP2 in this mouse may reflect the presence of a CA1-specific off-target site that was detected in one of the three additional injected mice that were tested for RNA editing by Sanger analysis (Table 1). The bystander off-target site was located 3 nucleotides 3’ to the target adenosine, and recoded the codon from lysine to arginine. Because this change occurred in the DNA binding domain, it could potentially prevent binding of MeCP2 to chromatin, although it is not a mutation causal to Rett syndrome [35]. Perhaps the off-target editing event in CA1 detected by Sanger sequencing and the lack of MeCP2 enrichment in the heterochromatin of CA1 neurons were both consequences of more virus being delivered to the CA1 region of those mice due to inconsistencies in the injection placement. Irrespective, because the off-target site was within the guide region, in future constructs, the aberrant editing can be eliminated by an A:G mismatch [19,42].

## Discussion

Both genomic and RNA base editing are potentially viable therapeutic approaches for treating human disease [20,43,44]. For example, genomic editing has been utilized to successfully repair a gene responsible for hearing loss in mice [45]. We have focused on RNA base editing because it does not have the same sequence constraints as DNA base editing. Additionally, as we have shown here, the rates of off-target editing, if it occurs, is usually graded with programmable RNA editing (Figure 3, [17,46]), in contrast to 100% mutation with genomic editing. Earlier work has demonstrated the potential of site-directed RNA editing for treating human disease. For example, heterologous expression in *Xenopus* oocytes of an Editase system repaired a chloride channel mutation underlying cystic fibrosis [15]. More recently, recoding of a pathogenic mutation in PINK1, associated with Parkinson’s disease, was achieved by full-length ADAR2-mediated editing in HeLa cells [47] and A-to-I and C-to-U recoding of 11 endogenous genes was achieved using an ADAR2-Cas13-guided system in HEK-293 cells [17,48]. The first demonstration of *in vivo* RNA editing was published recently using G>A mutant mouse models of Duchenne muscular dystrophy and ornithine transcarbamylase deficiency [21]. Our study herein is the first example of *in vivo* programmable RNA editing of a patient mutation in mice representing a neurological disease.

It is well accepted from analyses in cell lines and *in vivo* [17,21,46], and as we have shown in this study, that targeted RNA editing with exogenous hyperactive ADAR2 results in off-target editing events. Our case is likely the worst case scenario because we injected directly into hippocampus a very high titer of virus, and systemic injections, required for behavioral analysis in Rett mouse models, will result in much lower expression of Editase/cell. Several groups are already beginning to identify Editase molecules with higher specificity and efficiency [17,49], which will need to be tested *in vivo*, and the Rett mouse models will again be ideal for determining phenotypes. Importantly, despite the off-target events, in the mice expressing Editase and *Mecp2* targeting guide, we observed mean editing rates of 50% at the target adenosines in *Mecp2* RNA, in three different hippocampal neuronal fields containing multiple neuronal types [50-54] (Figure 2b). Further, the amount of MeCP2 associated with heterochromatin in these mice was also increased to 50% of wild-type (Figure 4d-f). One question that arises is how much MeCP2/cell is required to effect a behavioral improvement in symptoms? A mouse model containing a hypomorphic allele that reduces *Mecp2* RNA levels to 55% of wild-type levels does not exhibit overt Rett-like symptoms [55]. This result suggests encouragingly that the editing rates we achieved by programmable RNA editing could result in significant symptom improvement or stabilization of Rett symptoms if enough cells were repaired to this level. Related to this issue, the uniformity of editing and association with heterochromatin in the “repaired” mice also suggests that programmable editing rates may not range significantly across neuronal subtypes in mice, indicating that this level of repair may be achievable throughout the brain following systemic expression of the Editase system. Interestingly, a recent study also found that native editing rates of 5HT_2_C transcripts within human cohorts were similar across at least two different brain regions [56].

The on-target editing rates determined by Sanger and whole transcriptome sequencing in neurons in this study were consistent with the amount of heterochromatin-associated MeCP2 estimated from our imaging experiments (compare Figure 4d-f and Figure 2b). While the presence of immuno-labeled MeCP2 in heterochromatin is an indirect measurement of MeCP2 DNA binding ability, many previous studies have indicated that it is an excellent proxy based on comparisons with *in vitro* binding studies [40,41,57]. We noted that the variability of MeCP2 heterochromatin association in mutant neurons in mice infected with the *Mecp2* targeting guide was significantly less than that of native MeCP2 in wild-type neurons (dentate gyrus, p<0.0001; CA1, p=0.0025; CA3 p<0.0001). This result could reflect an as yet unidentified compensatory mechanism in the heterochromatin in mutants, from loss of MeCP2 from inception, which reduces the accessible number of MeCP2 binding sites. Further studies examining programmable Editase:guide editing at earlier time points may shed light on this intriguing observation.

## Acknowledgments

The authors thank Christine-Schmidt-Weber and Justine Nguyen for excellent technical help with mouse husbandry. We thank Joshua Rosenthal (Marine Biological Laboratories, MA) for his help and advice in applying the Editase system to Rett syndrome, and James M. Wilson, (University of Pennsylvania) for the p5E18-VD2/9 plasmid. We also appreciate discussions and encouragement from the laboratories of Adrian Bird (Wellcome Trust Centre for Cell Biology, University of Edinburgh), Michael E. Greenberg (Harvard Medical School), and Paul Brehm (Vollum Institute, Oregon Health and Science University). The work was funded by an NIH Director’s Transformative Research award (NS087726) to G.M., Joshua Rosenthal, Paul Brehm and John Adelman and NIH R01 NS088399 to H.N. Grants from the Rett Syndrome Research Trust to G.M. and J.R.S. are also gratefully acknowledged.

## Author contributions

J.R.S and G.M designed the research. J.R.S., S.Y.K., J.F., Z.S. performed the research. H.N. contributed AAV advice, reagents and analytic tools. S.J., S.K.M and performed the bioinformatics analysis with suggestions from J.R.S. J.S., S.Y.K., S.J., S.K.M and G.M. analyzed data and J.R.S and G.M. wrote the paper.

## Competing interests

G.M. is a co-founder of Vico Therapeutics. She holds equity but is not an employee and her laboratory receives no funding from Vico. The other authors declare no competing interests.

## Data availability

RNA-seq and exome sequencing data files have been uploaded to the Sequencing Read Archive (SUB7012760).

## Methods

### Animal Use

All animal procedures were approved by Oregon Health and Science University Institutional Animal Care and Use Committee. Mice were housed with littermates on a 12:12 light/dark cycle. The generation of *Mecp2*^*317G>*A^ mice and genotyping protocols have been described previously [19].

### Plasmid constructs

The creation of the plasmids containing the AAV vector genomes that were used for AAV vector production was described previously [19]. Guide sequences are shown in Supplementary Table 1. Plasmid DNA to generate viral vectors was prepared using the NucleoBond Xtra Maxi endotoxin free kit (Takara Bio) prior to production. The AAV helper plasmid expressing the AAV2 Rep proteins and the AAV-PHP.B capsid protein (i.e., the AAV-PHP.B helper plasmid) is a derivative of the AAV9 helper plasmid, p5E18-VD2/9 [58] and was constructed by inserting a PHP.B 7-mer peptide-coding DNA sequence [59] into the wild-type AAV9 capsid protein open-reading frame between the amino acid positions 588 and 589.

### AAV vectors

AAV vectors used in the study were produced in human embryonic kidney HEK 293 cells (AAV-293, Agilent, RRID: CVCL_6871) by an adenovirus-free plasmid transfection method and purified by two rounds of cesium chloride (CsCl) density-gradient ultracentrifugation followed by dialysis as described elsewhere [60]. To package AAV vector genome in the AAV.PHP.B capsids, we used the AAV-PHP.B helper plasmid as described above. The purified AAV vectors were in PBS with 5% sorbitol (w/v) and 0.001% Pluronic F-68 (v/v). The titer of each AAV vector was determined using quantitative dot blot using a probe generated against the Editase-coding sequence.

### Stereotaxic injections

P28-P35 *Mecp2*^*317G>A*^ male mice were deeply anesthetized with 4% isoflurane (vol/vol) and stabilized in a custom stereotaxic apparatus, modified from a David Kopf system. After being placed in the apparatus, mice were kept under 2% isofluorane (vol/vol) for the remainder of the surgery. A dental drill was used to make holes in the skull and each hippocampal hemisphere was injected using a pulled glass micropipette (diameter 10-15 μm) backfilled with AAV. Injections were made at the following coordinates relative to Bregma: medial-lateral (ML): 1.40 mm, anterior-posterior (AP): 1.50 mm at depths of 1.50 and 1.65 mm; ML: 1.75 mm and AP: 2.25 mm at depths of 1.75 mm and 2.00 mm. At each location 2.75 x 10^9^ viral genomes of virus were delivered. Following injections animals were allowed to recover on a heated pad prior to being returned to their home cage.

### Laser capture microdissection

Three weeks after stereotaxic injection, *Mecp2*^*317G>A*^ male mice were anesthetized by intraperitoneal injection of 2,2,2-tribromoethanol and sacrificed by decapitation. Whole brains were washed in ice cold phosphate buffered saline (PBS), embedded in Tissue Freezing Medium (Electron Microscopy Sciences) and stored at -80° C. Sagittal sections (12 μm) were cut at -25° C using a cryostat and loaded on poly (L) lysine coated PEN 1.0 membrane slides (Zeiss, cat #415190-9041-000). Immediately after sectioning, slides were fixed in 70% ethanol, stained with an abbreviated hematoxylin staining protocol, and stored at -80° C. Pyramidal cells from the CA1 and CA3 regions of the hippocampus along with dentate granule neurons were isolated for RNA analysis and cerebellar tissue was isolated for whole-exome sequencing using the Zeiss Palm Microbeam system.

### Sanger sequencing analysis

Whole hippocampal tissue and laser captured hippocampal fields were isolated from male mice three weeks post stereotaxic injection. Total RNA from whole hippocampal tissue was isolated using Trizol reagent (Thermo Fisher Scientific) according to the manufacturer’s instructions. RNA was isolated from laser captured cell populations using the RNeasy Micro kit (Qiagen) according to the manufacturer’s instructions. All samples were tested for RNA purity using a Bioanalyzer 2100 and had integrity scores of > 8.5. RNA was reverse transcribed using the SuperScript III First-Strand Synthesis System (Life Technologies) and primed using oligo dT. Endogenous *Mecp2* cDNA was amplified for analysis using a forward primer in the 5’ untranslated region and a reverse primer located in the 3’ untranslated region PCR products were fractionated on a 1% agarose gel and purified using the QIAquick gel extraction kit (Qiagen) before being submitted for Sanger sequence analysis. Sanger sequencing was performed using an Applied Biosystems 3730xl 96-capillary DNA analyzer. All primers are listed in supplementary Table 1. The C/T peak heights of the antisense strand were quantified from the resulting four-dye-trace sequences using the Bioedit software package (www.mbio.ncsu.edu/BioEdit/bioedit.html) as previously described [19,38]. Quantification of editing rates was performed using the antisense strand because A/G peaks have more inconsistent heights [61]. All chromatographs in the figures are shown as the reverse complement to show the mixed peaks at the target adenosine.

### Whole transcriptomic analysis

cDNA libraries were made by the OHSU Massively Parallel Sequencing Shared Resource using the SMARTer RNA kit (Clontech). Library quality was assessed using a TapeStation 220 and libraries were quantified by qPCR using a KAPA Library Quantification kit. Libraries were sequenced using 100-cycle paired-end runs on a HiSeq 2500. Whole genomic DNA was isolated from laser captured samples using a PureLink genomic DNA isolation kit (Ambion) and quantified using a TapeStation 220. Libraries were made from 50 ng of genomic DNA using a Seq-Cap Exome Plus capture kit and quantified by qPCR using a KAPA Library Quantification kit. Exome libraries were then sequenced using 100-cycle single-read runs on a HiSeq 2500.

Whole exome DNA sequencing results were aligned to the C57/Bl6J reference genome using Bwa-mem 0.717. The RNA-seq results were aligned to the mm10 reference genome using Bowtie 1.2.2. Single nucleotide polymorphisms (SNPs) were defined as DNA sample calls, which had < 99% of the reads aligning to the reference nucleotide across all sequenced samples and were excluded from downstream analysis.

RNA editing events were identified by comparing the adenosine or thymine nucleotides from the reference DNA sequence to the RNA sequencing results using the REDItoolDnaRna workflow [62] with the following parameters:

-e,E exclude multihits for RNA-Seq, DNA-Seq
-d, D exclude duplicates for RNA-Seq, DNA-Seq
-p User pair concordant reads only (for RNA-Seq only)
-u, U Consider mapping quality for RNA-Seq, DNA-Seq
-m 20,20 Minimum mapping quality score for RNA-Seq, DNA-Seq
-a,A6-0 Trim 6 bases up and 0 bases down per read for RNA-Seq, DNA-Seq
-I,L Remove substitutions in homopolymeric regions
-v1 Minimum number of reads supporting variation
-n 0.0 Minimum editing frequency for RNA-Seq, DNA-Seq

To be considered for further analysis, an editing event had to be present in all three biological replicates from each sample type (non-targeting guide and targeting guide) but not present in the non-injected controls.

### Immunostaining

Mice were anesthetized by intraperitoneal injection of 2,2,2-tribromoethanol and sacrificed by transcardial perfusion of PBS, followed by 4% depolymerized paraformaldehyde. Brains were removed and equilibrated in 30% sucrose overnight at 4°C before being embedded in Tissue Freezing Medium (Electron Microscopy Sciences) and stored at -80°C. Sagittal sections (14 µm) were cut at -20°C using a cryostat and stored at -20°C. Sections underwent antigen retrieval with - 20°C acetone for 8 minutes followed by washes with water and PBS before treatment with boiling citrate buffer (10 mM sodium citrate, 0.1% Tween-20, pH 6.0) for 10 minutes. Sections were cooled to room temperature (RT) and incubated in blocking buffer containing PBST (0.01% Triton X-100 in PBS, pH 7.4) and 10% normal donkey serum (Jackson Immunoresearch labs) for 30 minutes at RT. Sections were incubated overnight at 4°C with rabbit anti-MeCP2 (rabbit mab D4F3, Cell Signaling, 1:500) and rat anti-HA (rat mab 3F10; Roche, 1:200) antibodies diluted in blocking buffer. Sections were washed 3x with PBST and incubated for 1 hour at RT with Alexa Fluor secondary antibodies (Thermo Fisher Scientific, donkey anti-rat 488 and donkey anti-rabbit 647, 1:500) diluted in blocking buffer. Sections were washed 3x with PBST, then washed again with PBS before being incubated with 300 nM 4′,6-diamidino-2-phenylindole (DAPI) in PBS for 20 minutes. After a final wash in PBS, sections were mounted using ProLong Gold antifade reagent (Thermo Fisher Scientific), which was allowed to cure overnight.

### Image Acquisition and Analysis

A Zeiss 710 laser scanning confocal microscope equipped with a 63x Plan-Apo objective was used to acquire sequential 1 µm optical sections for creation of Z-stack images. The field size corresponded to 18211.5 um^2^ with a resolution of 1024×1024 pixels. Fluorescence images corresponding to HA label (488 laser), MeCP2 label (633 nm laser) and DAPI (405 nm laser) were sequentially acquired. For each, the laser strength was set to sub-saturating levels corresponding to 0 to 255. These acquisition settings were then applied to all samples.

The fraction of HA immuno-labeled cells was determined by counting the fraction of DAPI positive nuclei that were also HA positive. Cells were determined to be HA positive if the fluorescence intensity was above non-injected controls. At least 100 cells were counted in each region of the hippocampus (CA1, CA3 and dentate gyrus) for each mouse using the ImageJ cell counter plugin (National Institutes of Health, https://imagej/nih.govij, version 1.60_65 (32bit)) and two *Mecp2*^*317G>A*^ mice were analyzed for each viral condition. As a proxy for the ability of MeCP2 to bind DNA, the MeCP2 intensity in pericentromeric heterochromatic foci was determined. Once again, the laser strengths for MeCP2 and DAPI were adjusted individually to fall within a nonsaturating 0-255 range and the acquisition parameters were held constant for all subsequent measurements. Regions of heterochromatin selected for measurement were based on both size (≥ 0.3 μm^2^) and average intensity (≥ 80 on a scale of 0 to 255). Using Image J, the corresponding MeCP2 fluorescence intensity for 4 to 6 heterochromatic foci per cell was determined on the basis of 4 x 4 pixel ROIs. These values were then used to generate the average intensity value for each cell. For each of two animals, 20 cell averages were generated for each of 3 regions of the hippocampus (CA1, CA3 and dentate gyrus). This process was repeated for both viral conditions (wild-type non-injected, *Mecp2*^*317G>A*^ targeting guide, *Mecp2*^*317G>A*^ non-targeting guide). The resultant amplitude distributions (n = 40 cells per condition) were subject to statistical comparison using a Kruskal-Wallis test followed by a Dunn’s multiple comparisons test. Further, for each hippocampal region the variance of MeCP2 fluorescence intensity for each condition was subject to statistical testing using an F-test for equality of variances.

### Statistical Analysis

All statistical tests with the exception of the whole transcriptome analysis were performed using GraphPad version 6.0 software (Prism). The rate of A-to-I editing from whole hippocampal samples was compared between viral conditions using an unpaired *t*-test and within isolated neuronal populations using one-way ANOVA followed by post-hoc Tukey’s post-hoc multiple comparisons tests. Western blots comparing the level of Editase protein and the number of RNA-seq reads aligning to the Editase coding sequence were compared using unpaired two-tailed *t-*tests. The level of MeCP2 fluorescence in heterochromatic foci was compared within hippocampal regions using a Kruskal-Wallis test followed by a Dunn’s post-hoc multiple comparisons test. The variance of MeCP2 fluorescence intensity was subject to testing using an F-test for equality of variances. All experimental results are expressed as the mean ± the standard deviation.

---

**Supplementary Figure 1:**
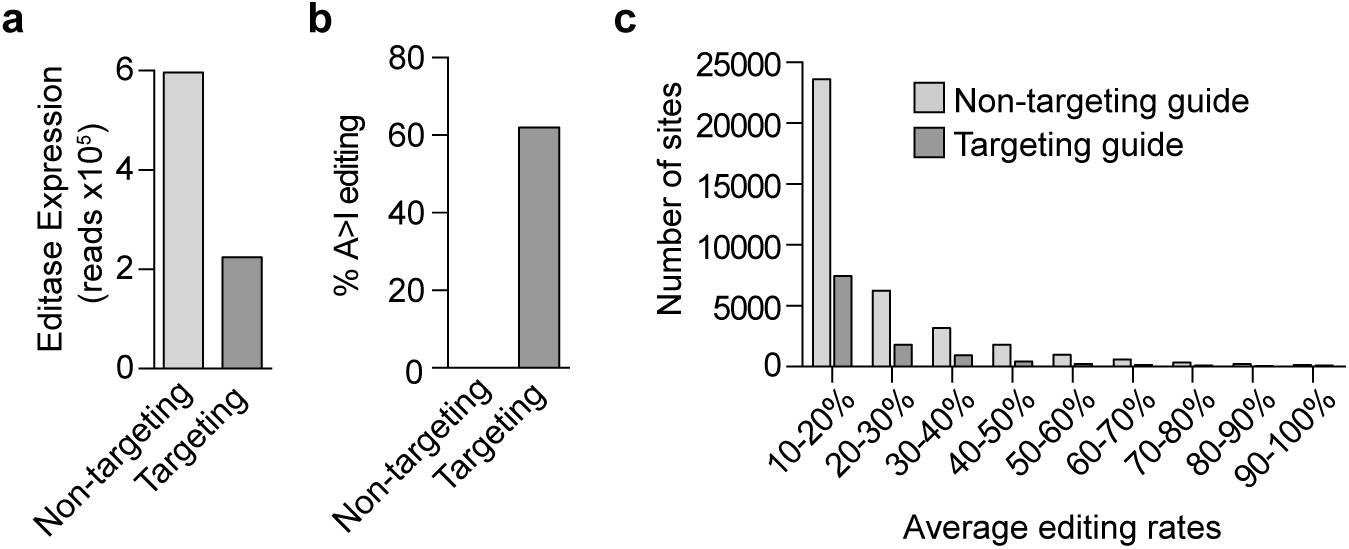
Whole transcriptomic RNA seq analysis of the same RNA used in our published study [19] of *Mecp2*^*317G>A*^ hippocampal neurons (DIV14), 7 days following transduction with AAV1/2 virus. Promoters, Editase and guide sequences are identical to that shown in Figure 1a in this study. a) The number of RNA-seq reads that aligned to the Editase coding sequence for each viral condition. b) Histogram showing that on-target editing is guide-dependent. One sample each condition. c) Histogram showing the number of off-target editing sites, binned according to the editing rates.

**Supplementary Figure 2:**
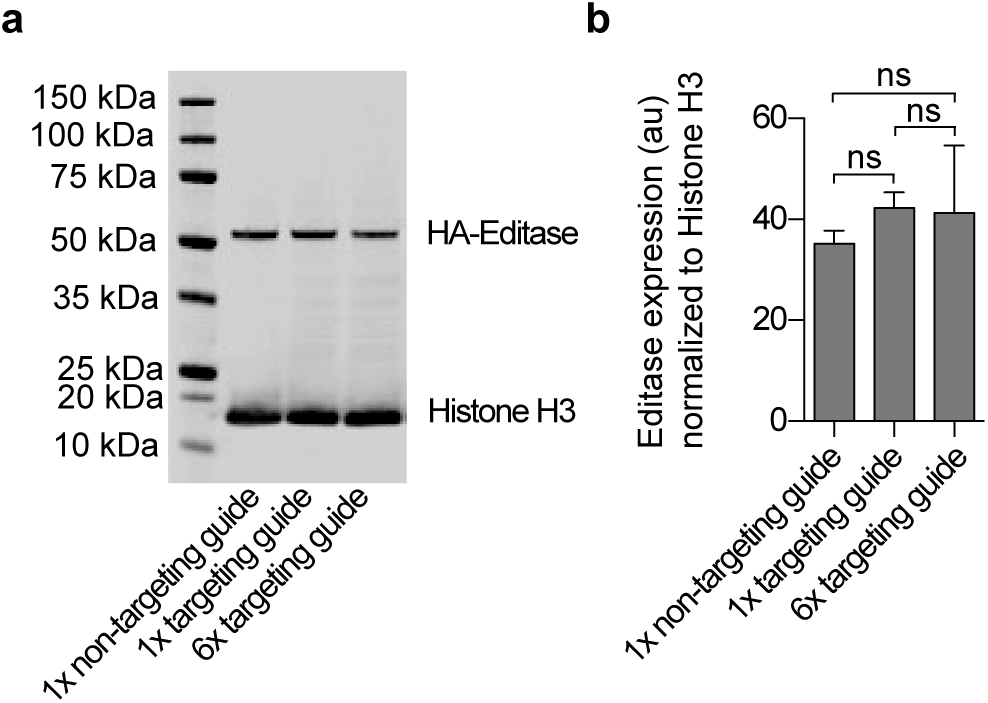
The number of copies of the human U6 promoter driving guide expression does not influence Editase protein expression. a), Representative immunoblot of whole cell lysates prepared from Neuro-2A neuroblastoma cells 72 hours after transfection with plasmids containing the Editase cDNA expressed from the human *Synapsin I* promoter and indicated guides. Blots were probed with anti-HA antibody for Editase detection and anti Histone H3 for loading control. b) Quantification of Editase expression from immunoblots (mean ± SD, n = 3 biological replicates), each condition normalized to Histone H3. au, arbitrary units. ns, not significant by One-way ANOVA and Tukey’s multiple comparisons test.

**Supplementary Table 1:**
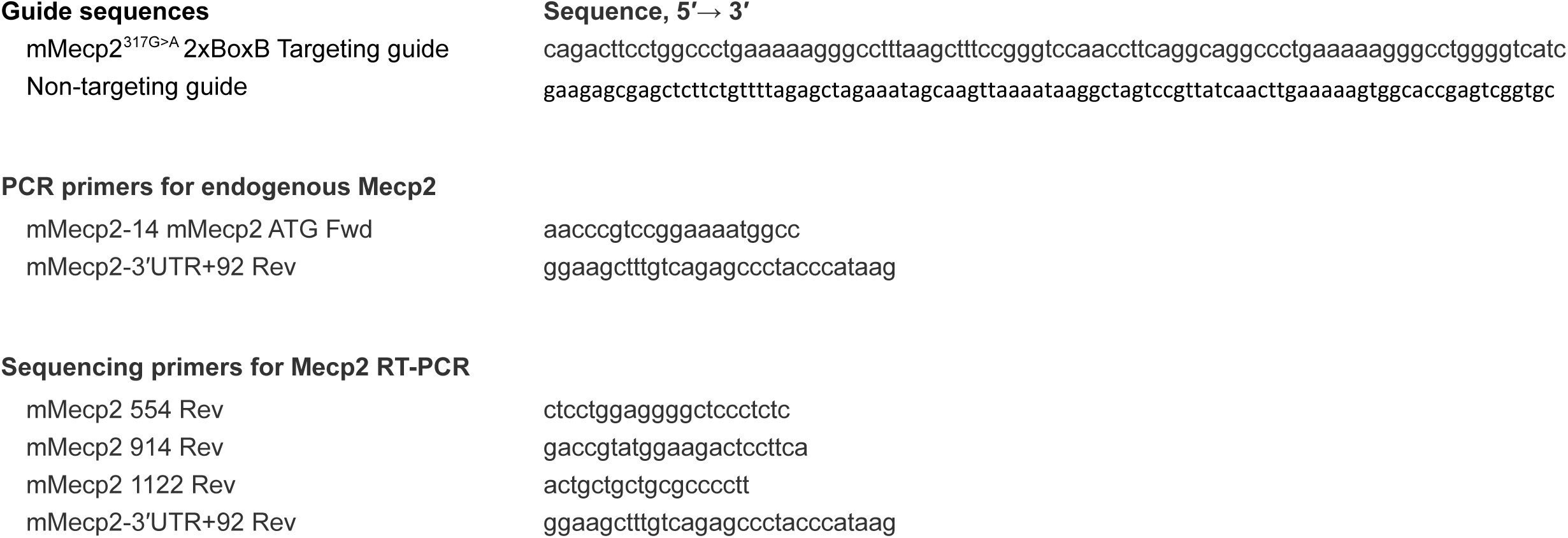
Guide sequences, PCR and sequencing primers

## Supplementary Methods

### Transient Transfections and Cell Culture

Neuro2A cells (ATCC CCL-131) were maintained in DMEM (Thermo Fisher Technologies) in 10% FBS (HyClone) at 37 °C in a 5% CO_2_ humidified incubator. For analysis of Editase protein expression, cells were seeded at a density of 1.25×10^5^ cells per well of a 12-well plate. After 24 h, cells were transfected with 1 µg of plasmid DNA containing the human *Synapsin I* promoter expressing Editase and six copies of the *Mecp2* targeting guide (pGM1258), one copy of the *Mecp2* targeting guide (pGM1267) or one copy of the non-targeting guide (previously referred to as Editase alone, pGM1186). Plasmid DNA was transfected using a 2:1 ratio of Lipofectamine 2000 (Thermo Fisher Scientific) and DNA in Opti-MEM reduced serum media (Thermo Fisher Scientific).

### Western Blotting

Transfected Neuro2A cells were lysed 72 hr after transfection using 100 µL of whole-cell lysis buffer (25 mM Tris, pH 7.6, 150 mM NaCl, 1% Igepal CA-630; Sigma, 1% deoxycholate, 0.1% SDS, protease inhibitor (Complete EDTA-free; Roche), 1 mM beta-mercaptoethanol, and 250 units per mL benzonase (Sigma-Aldrich)). Lysates were centrifuged at 16,000 × *g* for 10 min at 4° C and the soluble fraction isolated. Protein concentrations from the soluble fraction were measured using the BCA protein assay kit (Pierce Biotechnology). 20 µg of protein lysate per sample was separated on NuPage 4–12% Bis–Tris gels (Thermo Fisher Scientific) in Mops-SDS running buffer (Thermo Fisher Scientific), and transferred onto a nitrocellulose membrane (GE Healthcare Life Sciences). Membranes were blocked with 3% BSA in 1× TBST (TBS with 0.05% Tween 20) for 1 h, then incubated with mouse anti-HA (Biolegend; 1:1000) and rabbit anti–Histone H3 (Cell Signaling; 1:5000) overnight at 4 °C. After washing three times with 1× TBST, blots were incubated with anti-mouse IgG DyLight 680 (1:10,000 dilution; Thermo Scientific) and anti-rabbit IgG Dylight 800 (1:10,000 dilution; Thermo Scientific) diluted in 3% BSA in 1xTBST for 1 h. Blots were imaged and quantified using the Odyssey Imaging System (LI-COR Biosciences).

